# Influence of membrane-cortex linkers on the extrusion of membrane tubes

**DOI:** 10.1101/2020.07.28.224741

**Authors:** Alexandru Paraschiv, Thibaut J. Lagny, Christian Vanhille Campos, Evelyne Coudrier, Patricia Bassereau, Anđela Šarić

**Author notes:** Electronic address (Anđela Šarić).

## Abstract

The cell membrane is an inhomogeneous system composed of phospholipids, sterols and proteins that can be directly attached to underlying cytoskeleton. The linkers between the membrane and the cytoskeleton are believed to have a profound effect on the mechanical properties of the cell membrane and its ability to reshape. Here we investigate the role of membrane-cortex linkers on the extrusion of membrane tubes using computer simulations and experiments. In simulations we find that the force for tube extrusion has a non-linear dependence on the density of membrane-cortex attachments: at a wide range of low and intermediate densities of linkers the force is not significantly influenced by the presence of membrane linking proteins and resembles that of the bare membrane. For large concentrations of linkers however the force substantially increases compared to the bare membrane. In both cases the linkers provided membrane tubes with increased stability against coalescence. We then pulled tubes from HEK cells using optical-tweezers for varying expression levels of the membrane-cortex attachment protein Ezrin. In line with simulations, we observed that overexpression of Ezrin led to an increased extrusion force, while Ezrin depletion had negligible effect on the force. Our results shed light on the importance of local effects in membrane reshaping at the nanoscopic scales.

## I. INTRODUCTION

Biological membranes can dynamically change their shape as a response to an external mechanical stress. The application of a localised force to a membrane leads to the formation of membrane tubes or tethers which are thin elongated cylindrical structures [1]. Membrane tubes are involved in a wide variety of cellular processes such as motility, mechanosensing, signalling, inter-cellular and intra-cellular trafficking [2–4]. Tubes can be artificially extracted through hydrodynamic flow experiments, atomic-force microscopy, optical or magnetic traps, both from reconstituted vesicles as well as from live cells [5–9]. In the case of giant vesicles and bare membranes the force-extension curve of the extrusion process has been well characterised both in theoretical models and in experiments [10–15]. The shape of the tube and characteristic extrusion force were used to directly infer the mechanical properties of the membrane, where the force was found to depend on the membrane bending rigidity *κ* and surface tension *γ* as 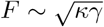, while the radius of the tube scaled as 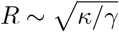.

Plasma membrane of eukaryotic cells is connected to the underlying cytoskeleton via linking proteins such as ERM (Ezrin, Radixin and Moesin) and myosin 1 proteins [16, 17]. The influence of these linkers on the force required to extrude a tube is still a source of discussion. The attachment to the actomyosin skeleton has been suggested to play a role in the extension of membrane tethers, by providing resistance to the flow of the membrane towards the tube [18]. It was also shown that reducing the attachment of the plasma membrane to the actomyosin cortex, via myosin 1b or Ezrin depletion, reduces the static force required to extrude a tether in zebrafish progenitor cells [19]. A similar result was obtained with myosin 1g in lymphocytes [20]. Furthermore, Ezrin phosphorylation that increases membrane-cortex attachment was shown to increase the tether force in lymphoblasts [21].

Taken together, this data has been interpreted as evidence that membrane-cortex attachments effectively change the membrane tension. In this interpretation the tension is increased by the strength of the membrane-cortex adhesion per unit area, which in turn also increases the force needed to extrude a membrane protrusion [22]. Alternative pictures have also been put forward, in which the membrane flows around the membrane-cytoskeleton attachments into the tether, such that the attachments do not contribute to the extrusion force, resulting in a force that resembles that of the bare membrane [23, 24].

Here we use coarse-grained computer simulations to study the role of the cytosleketon-membrane attachments on the extrusion of membrane tubes in isolation from other effects that might be present in cells. Our simulations do not explicitly assume driving forces, but only assume the ingredients – a simple model of a fluid membrane that is sparsely attached to the underlying surface. We then carry out computational experiments where we extrude protrusions and measure the membrane response for a wide range of membrane-cortex attachment densities. We complement our observations with optical-tweezers-based tether pulling on HEK293T cells adhering on a substrate to measure the impact of the ERM protein Ezrin on membrane mechanics.

Our results indicate that the two previously polarised views can in fact co-exist: we find that at low membrane-cortex linker densities the extrusion force is not influenced by the cortex attachments and is equal to that of the bare membrane, while at high attachment densities the force strongly increases compared to the bare membrane. At low linker densities the membrane flows around the linkers and the linkers avoid entering the tube to minimise the energetic cost of extrusion, hence the membrane-cortex attachment does not contribute to the force. At high linker densities, however, the linkers cannot avoid entering the tube composition any more and necessarily start detaching, causing the substantial increase in the extrusion force accompanied by a slow tube relaxation dynamics. In agreement with the simulations findings, our experiments show that for cells that display moderate level of cortex-linker attachments, depleting ERM protein Ezrin has negligible influence on the extrusion force but impedes the tube relaxation, while overexpressing Ezrin leads to a significant increase in the force. Our results point to the importance of local effects – the redistribution of the membrane-cortex attachments around the protrusion – when mechanically probing cellular membranes at the nanoscale.

## II. RESULTS

### The force profile for the bare membrane

We use a Monte Carlo model in which the fluid membrane is described as a dynamically triangulated elastic sheet, while the cytoskeletal linkers are modelled by attaching some of the vertices to the surface underneath it by dynamic finite harmonic springs. This model reproduces well the mechanical properties of biological membranes, such as bending rigidity and fluidity, but does not take into account the explicit bilayer structure and the details of the membrane and protein structures. We also repeated our key results using a self-assembled one-particle-thick membrane model within molecular dynamics simulations, as discussed in SI.

We first make sure that our model reproduces previously measured force-extension profiles for non-attached membranes. To that end we apply a vertical pulling force on a small membrane patch in the membrane centre. This leads, at first, to the extrusion of a cone-like structure (Fig. 1a). The force-extension profile shows a linear elastic response until a critical threshold is reached. The force then drops by around 8% and the protrusion undergoes a shape transition from the catenoid to a thin tubular structure as shown in Fig. 1b and the extrusion force eventually reaches a plateau. This extension profile is consistent with experimental and theoretical studies of tether extrusion on free membranes that show the familiar profile of a barrier followed by a plateau [10]. It has been shown that the height of the barrier is influenced by various factors such as membrane composition, speed of pulling and the area of contact between the membrane and the pulling object, while the plateau is typically determined only by the membrane bending rigidity and surface tension [12, 24–26]. We next explored how this force-extension curve is influenced by the addition of proteins linking the cytoskeleton to the membrane.

**FIG. 1:**
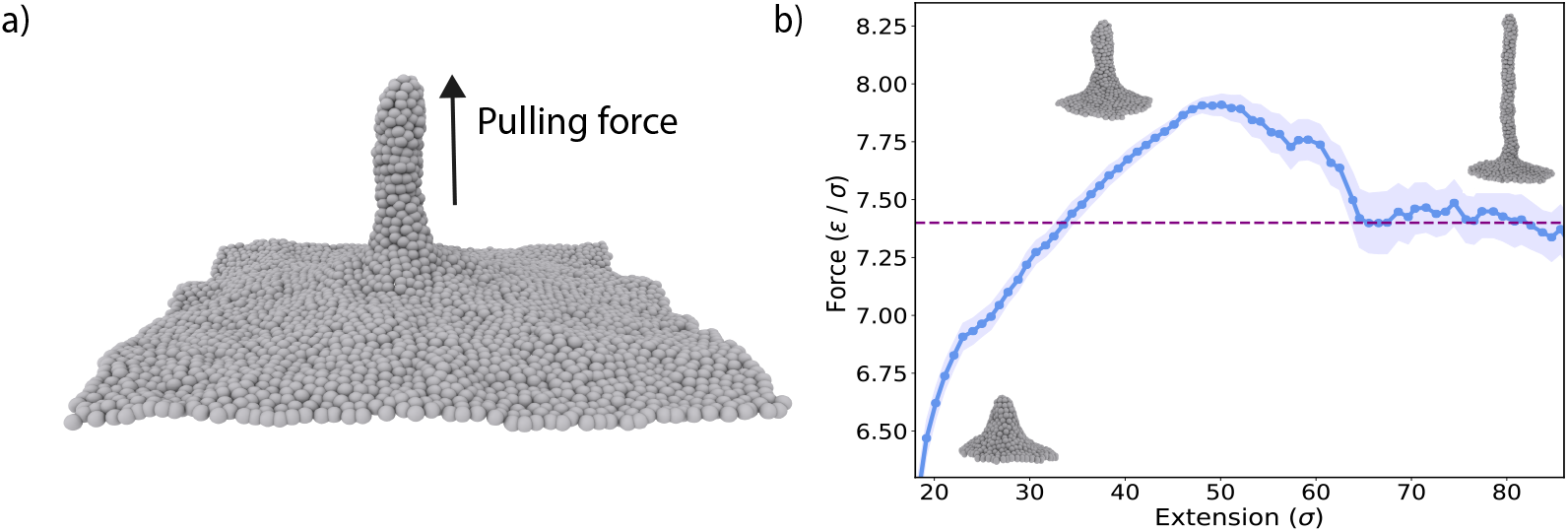
The membrane is extruded into a tubular structure. (a) The membrane is modelled as a uniform thin sheet made of a single layer of phospholipids (grey beads). The membrane is pinched at a central bead and a tubular structure is extruded. b) The force-extension curve for a homogeneous membrane (in the absence of any cortex-linkers) shows a peak at an extension of approximately *l* = 50*σ* which corresponds to a transition in the protrusion’s aspect from a truncated cone to a thin elongated tube. The curve eventually reaches a plateau after which the extrusion proceeds at the same force. The shaded area represents the standard error for measurements from 10 different simulation seeds.

### Extrusion force has a non-linear dependence on cortex-linker density

Cytoskeletal attachments, or linkers, were introduced into the system by connecting a fraction of membrane beads to a virtual plane beneath the membrane by a generic, finite harmonic, potential (Fig. 2a). Such a bond can be broken if sufficiently extended. Although some membrane-cortex linkers can exhibit more complex behaviour such as catch-bond [27], this simple generic form of binding is believed to represent well the ERM proteins [28].

**FIG. 2:**
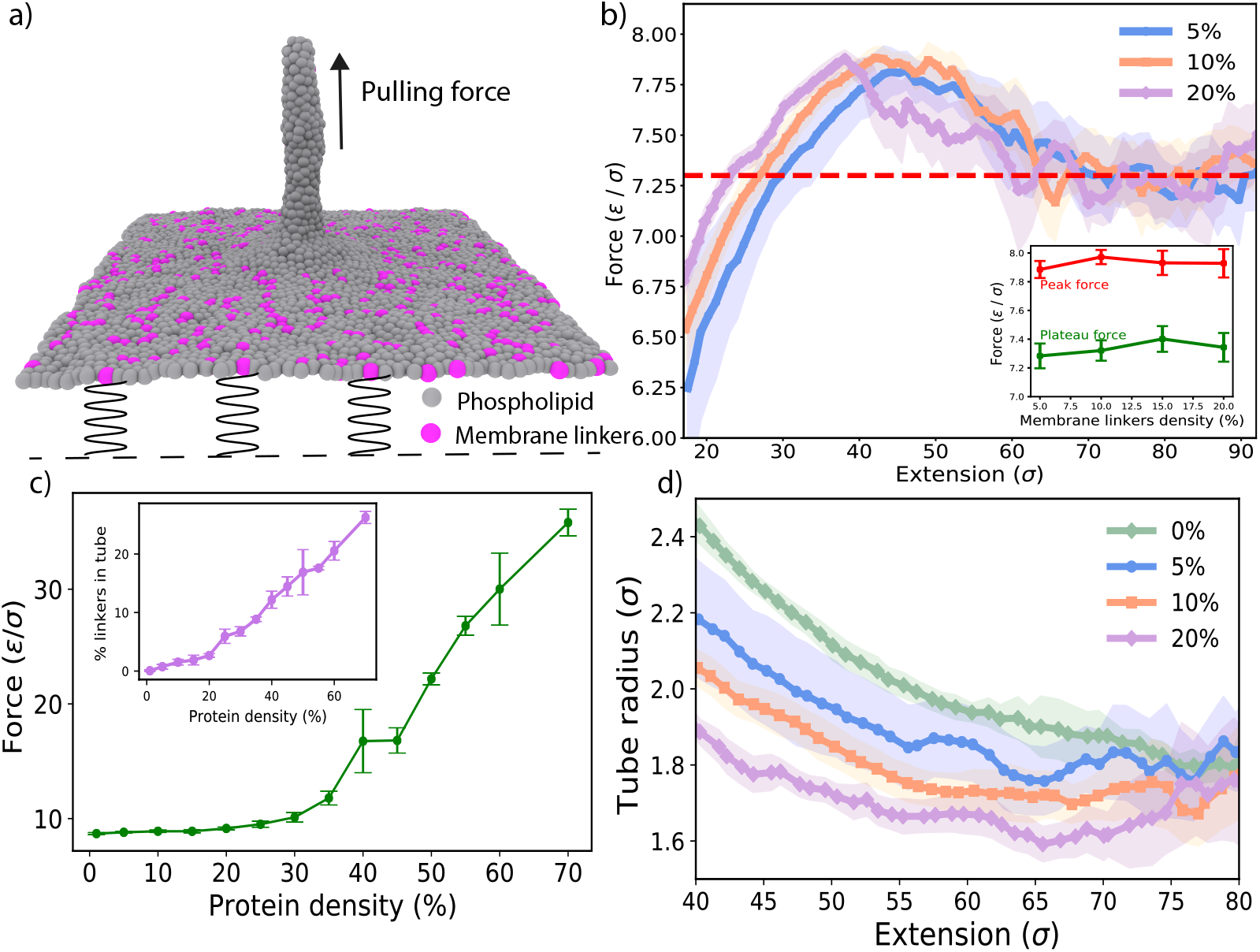
Force-extension curve in the presence of membrane linkers. (a) Side-view of a tubular structure pulled in the presence of membrane linkers (magenta beads). The linkers are restrained on the vertical plane with a harmonic potential. (b) Force extension curves at small and intermediate linker densities are not significantly affected by the presence of linkers. The peak and plateau forces remain constant independent of protein density (legend indicates the protein number density, *ε* is the simulation unit of energy of 1 *k*_*B*_*T* and *σ* is the simulation unit of length). c) The force on the tube tip measured for the target extension of 60*σ*, averaged between 1*·*10^6^ and 2 *·* 10^6^ MC steps. At high expression levels of the proteins the force increases substantially. Inset shows the fraction of proteins in the composition of the tube. The error bars and shaded regions in (b)-(d) represent the standard error of the measurements over 5 different simulation seeds. d) The tube radius decreases as the tube is extruded. The protein density modulates the radius.

As the membrane tube is extruded from such a pinned membrane, we observe that the linkers tend to disperse away from the tube since there is a high energetic cost associated with opposing the pulling force and entering the tether. This causes the force required for extrusion not to be significantly affected by the linker attachments for low protein fractions, as shown in Fig. 2b. In the early stages of the tube formation, the force required to reach the same extension is slightly greater at higher protein concentrations [19], but the plateau force does not show a significant difference at low to moderate linker concentrations. However, if the linker density is high enough, the growth of a protrusion requires a large scale concerted reorganisation of the linkers in order to facilitate the transport of sufficient material into the tubular structure. In this case the linkers cannot effectively exclude themselves from the tube and will experience a pulling force, at the same time slowing down the dynamics of tube extrusion.

The growth of a protrusion hence becomes expensive, resulting in an exponential increase in the pulling force for higher protein concentrations, as seen in Fig. 2c, eventually fully preventing tube extrusion. Taken together, our model shows that the extrusion force can have non-linear dependence on the linker densities – at low densities the force will not be influenced by the linkers, while at higher linker densities the force will substantially increase.

To check that our results are not influenced by the details of our model – namely the Monte Carlo routine that might not capture the correct dynamics – we repeated our results using a non-triangulated spherical membrane within a molecular dynamics (MD) framework. As shown in the SI Section II H and Fig.S10–12, our results remain qualitatively unchanged: the molecular dynamics model also shows a non-linear dependence of the force on the linker density (Fig. S12). The only differenc is that the critical linker density at which the force increases is shifted to lower values compared to the Monte Carlo model. These results indicate that the non-linear dependence of the force on the linker density is a robust result, which is not model dependent, but that the point of the crossover from the flat to the exponential increase will depend on the details of the model and the system – namely the interactions of the proteins with the cortex and the membrane fluidity.

Finally, we observe that presence of linkers also causes a slight reduction in the tube radius (Fig. 2d): as the linker density increases, there is less material freely available for the tube growth, resulting in a decrease of the tube diameter (see also Fig. S11 for the MD results).

### Linkers are excluded from the tube at intermediate densities

Fig. 3a further depicts how the linker proteins dynamically rearrange around the tube during tube extrusion in our simulations. To minimise the strain imposed on the membrane the linkers avoid the region around the tube neck, giving rise to a concentration gradient in proteins around the tube. To minimise the perturbation to the membrane the proteins also tend to segregate into small protein-rich clusters (Fig. 3b and Fig. S10). These results indicate that pulling the tube promotes the local aggregation of proteins by depleting the available phospholipid membrane area. The tube itself can contain small concentrations of trapped proteins (Fig. 3c). We observed that the partitioning of proteins between the tube and the membrane is influenced by the linker binding energy, with higher binding energies resulting in stronger linker exclusion from the tube and smaller tube radii (Fig. S4 and Fig. S5).

**FIG. 3:**
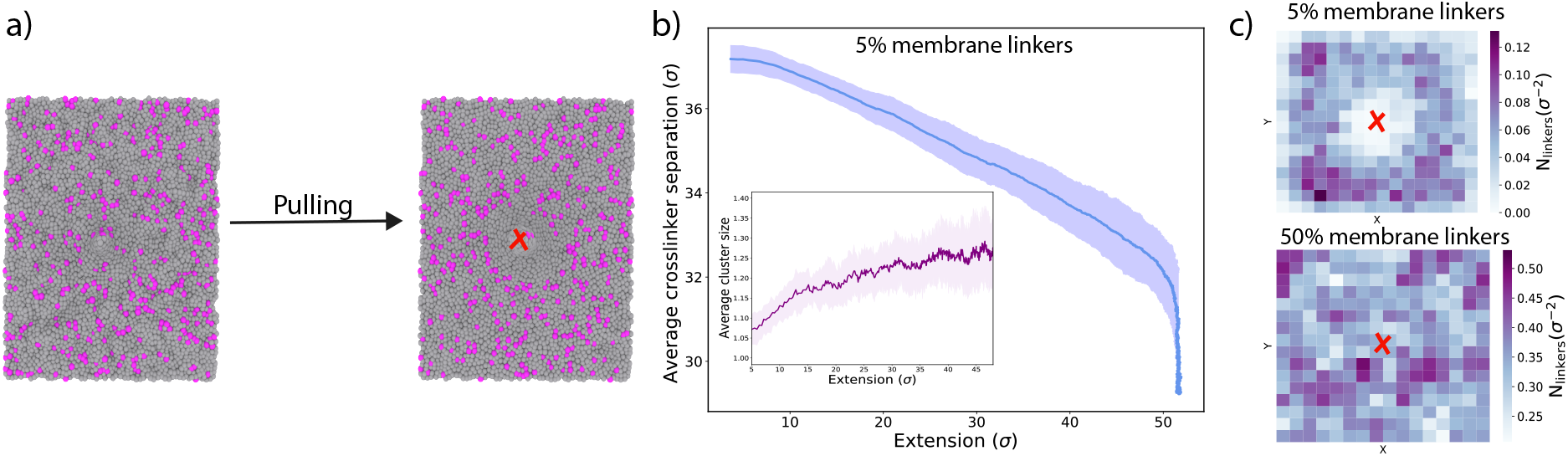
The linkers are excluded from the tube at intermediate linker densities. (a) Top-view of the membrane before and after the extrusion of the tube. The linkers avoid the tube and prefer to segregate into small clusters. (b) The average separation between the linkers decreases as the tube is extruded from the membrane. The linkers cluster size (dark purple line, inset) increases as the extrusion progresses (linkers are considered in the same cluster if the distance between them is less than 3*σ*) . The data is obtained from averaging over 20 simulations where the tube is extruded to a target length of *l*_0_ = 60*σ*. The shading represents the standard error of the measurements over 5 different simulation seeds. (c) Heatmaps showing the number of linkers per unit of membrane area at low (5%) and high (50%) linker densities. The squares are weighted according to the membrane area they contain. The central region, where the tube is extruded, (marked with a red ‘X’ symbol) is depleted of linkers at low densities, showing that proteins prefer to diffuse away from the tube. At high densities, the proteins are distributed uniformly with a significant fraction trapped inside the tube.

### Experiments: Influence of Ezrin expression on tube extrusion in cells

In order to qualitatively compare simulation results to experiments, we pulled tethers from adherent cells as depicted in Figure 4a with a typical length of 10 *µ*m. As previously shown, the initial force overshoot depends on the interaction area between the bead and the plasma membrane [12] and is not well controlled under experimental conditions (Figure 4a – step 3 and Fig. S16a). Upon tube formation the force next decreases and rapidly reaches a plateau. To measure the static tether force, we waited 10 seconds to let the force equilibrate (Figure 4a – step 4, and Figure S16a). In agreement with previous reports [29], we confirmed that the tether force does not depend on tether length over a broad range (Fig. S16b). In addition, special care was taken to ensure the measurement of forces for single tethers only.

**FIG. 4:**
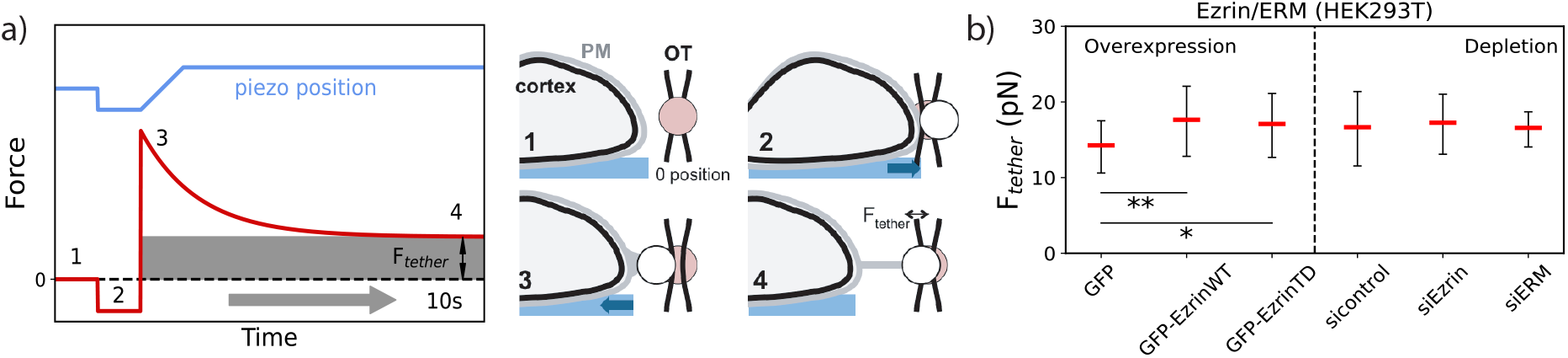
Measuring plateau force for varying Ezrin levels. (a) Force regimes observed during tether pulling from cells corresponding to the steps depicted on the right. A bead at equilibrium position (1) is brought into contact with a cell (2). When the stage is pulled away from the trap, an initial force peak required to detach the membrane from the underlying actin cortex (3) is quickly followed by a stable force *F*_*tether*_ (4). (b) Plateau forces in HEK293T cells overexpressing GFP (n=19), GFP-EzrinWT (n=28), GFP-EzrinTD (n=21) or treated with control siRNA (sicontrol, n=36), Ezrin siRNA (siEzrin, n=23), or ERM siRNA mix (siERM, n=18). mean*±*S.D(\*\**p* = 0.008; \**p* = 0.03).

We measured a tether force of 16.0 ± 2.7 pN in HEK293T cells, which is at the lower end of values reported for adherent cells [30] (Figure S14). We checked that the tether force is unchanged when the membrane receptor EphB2 is expressed (HEK-EphB2). It is similar in HeLa cells, yet higher in RPE-1 cells (Figure S14), highlighting that the static tether force varies depending on the cell type. Next we studied the effect of the manipulation of the expression level of ERM on tether force. Since the transfection process can perturb the cells, we compare the measurements after depletion or over-expression with control cells transfected according the same protocol, but without plasmid. GFP-Ezrin is binding actin in a phosphorylation-dependent manner, whereas the phosphomimetic mutant GFP-EzrinTD permanently links actin to the membrane [31] (Figure S15b).

We found that the depletion of Ezrin or ERM using siRNA [32] (Figure S15c) did not change the tether force compared to the control (Figure 4b), while the expression of recombinant Ezrin-WT and Ezrin-TD proteins significantly increases the force in HEK cells as compared to control (Figure 4b and Figure S15). These results favourably compare to our simulations where we found that the force to extrude tethers increases only when the expression level of linkers is substantially increased. Moreover, depleting Ezrin/ERM enabled faster tube relaxation (Fig. 5b), in agreement with what has been observed in simulations (Fig. 5a). These dynamic results show that the presence of linkers imposes a barrier for the diffusion of membrane components, effectively increasing the local friction [24, 33].

**FIG. 5:**
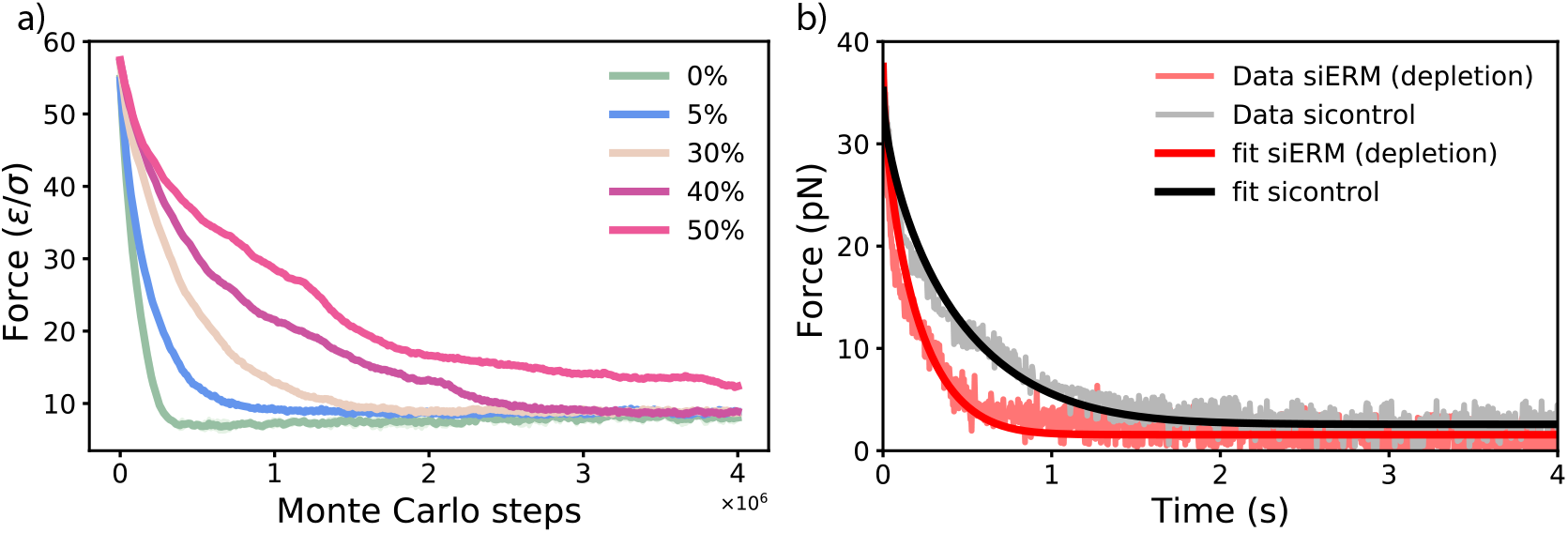
Tether extrusion dynamics depends on the density of linkers. a) Simulations: As the tube is extruded the force acting on the protrusion’s tip decays to a constant non-zero value. The dynamics of this relaxation is slower at higher linker density. Each curve represents an average over 5 simulation repeats. b) Experiment: upon fast tube elongation the force relaxes to a constant non-zero value with a time-scale that is faster upon depletion of ERM/Ezrin.

## Linkers stabilise single isolated membrane tubes

Membrane tubes pulled from bare membranes have been previously shown to merge in each other’s proximity, as the free energy change involved in this process is always favourable [10, 34]. To test the influence of the linkers on the interactions between protrusions we go back to our simulations. We pulled two tubes at an initial separation of Δ*L* = 20*σ* and allow them to diffuse freely. For bare membranes we observe that the two tubes grow and eventually merge through the basal truncated cone-like structure into a characteristic Y-shape as shown in Fig. 6a. This finding is in qualitative agreement with the structures observed in double tube experiments [34]. Interestingly, when the linkers are introduced, the probability of tube fusion decreased substantially and is lower at higher linker densities (Fig. 6c).

**FIG. 6:**
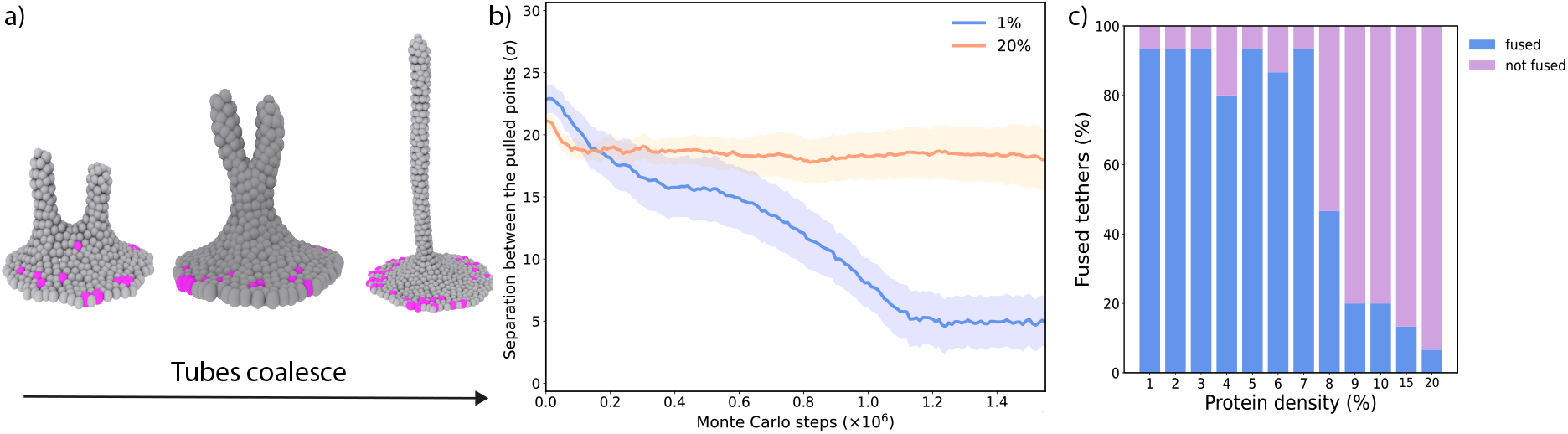
Protrusion coalescence is hindered by linkers. (a) Tubes coalesce into a Y-shaped structure. (b) The distance between the pulled points decays quickly at low protein densities, showing the tendency of adjacent tubes to coalesce. At higher densities, the linkers will maintain a stable distance, as they are prevented from fusion. The shading represents the standard error of the measurements over 20 different simulation seeds. (c) The percentage of tubes that fuse into a single structure decreases at greater linker densities.

To quantify this effect we measured the relative separation between the tips of the two pulled tubes during the course of the simulation (Fig. 6b). The average separation between the tubes decreases more slowly at greater protein densities, indicating that the linkers act as obstacles in the merging process. The fusion requires the neck of the tubes to be depleted of linkers before it can take place. The proteins thus act as a barrier for the fusion process, stabilising isolated tubes. Our findings are in an excellent agreement with the study of Nambiar et al [35], where the authors found the membrane tethers coalesce more frequently when the membrane-cortex attachment is decreased by depleting myosin 1 proteins, just like in our simulations. It can thus be concluded that the membrane linkers can serve to stabilise protrusive structures involved in physiological processes. Along the same lines, Datar et al. found that the membrane tethers move more easily on the surface of axons when the cytoskeleton is disrupted [33].

## III. DISCUSSION AND CONCLUSIONS

Using coarse-grained simulations, where lipid components and membrane-cortex attachments are modelled explicitly, we found that the force needed to extrude a protrusion has a non-linear dependence on the density of the membrane-cortex attachments. At low and moderate densities of the attachments the amplitude of the force needed to extrude a protrusion is unchanged compared to the bare membrane. In our model this is caused by i) the ability of the membrane to flow around the attachments, effectively excluding the attachments from the tube and ii) the ability of the attachments to diffuse away from the tube rather than to oppose the tube extrusion. In contrast, when the linker density is increased above a certain threshold, the force needed for tube extrusion increases significantly, as the linker exclusion out of the tube becomes prohibitively slow, as also demonstrated by a slower relaxation of the force to the equilibrium value.

The linkers’ diffusion in our model effectively captures dynamic ERM attachment/detachment from the cortex [36]. To test the influence of the linkers’ diffusion on our key results on the nonlinear dependence of the extrusion force on the linker density, we repeated these measurements for the case where the linkers cannot rearrange and diffuse in the membrane plane, but can still detach. As shown in Fig S9, the non-linear behaviour of the force versus linker density remains unchanged, although the point of cross-over was again shifted to lower the linkers’ density.

Our simulations indicate that great care should be taken when modelling the membrane-cortex attachments in an effective way, for instance by describing them as a uniform membrane-surface adhesion or rescaled membrane tension. Here we clearly show that the local effects, such as the exclusion of linkers from the tube, can have non-trivial effects on the resulting extrusion force, as well as on the protrusion radius.

Our experimental results align well with such non-linear effects observed in simulations and show that depletion of Ezrin/ERM does not have significant influence on the plateau force, while the over-expression does. Such a response could indicate that the density of linkers in HEK cells might lie close to the critical transition from a plateau to exponential part of the force versus linker density curve.

Previous papers based on static tether force measurements have reported effects of Ezrin and myosin 1 isoforms on effective membrane tension in non-adherent cells such as lymphocytes [20], or in cells with high blebbing propensity [37]. By interfering with the membrane-cortex attachment Diz-Muñoz et al. showed that a decrease in the attachment led to a decrease in the force required to pull a membrane tether from developing zebrafish embryo [19]. Recently, Berget et al. reported that incorporating a synthetic linker that increases the membrane-cortex attachment in embryonic mouse cells led to an increase in the tether-extrusion force [38]. Similarly, de Belly et al. [39] observed that mouse embryonic stem cells that display membrane blebbing, which is believed to be a sign of low membrane-to-cortex attachment, require lower force for tether extrusion than non-blebbing cells. In all of the above cases the forces required for tube extrusion from embryonic cells (∼50 pN) were substantially higher than in the case of HEK cells probed here (∼10 pN). It is hence likely that the expression level of membrane-cortex attachments in embryonic cells is naturally higher than in epithelial cells studied here, which would cause the experiments to probe the right-most part of the non-linear curve presented in Fig. 2c, where the force severely depends on the linker concentration.

he critical density of linkers after which the force increases is expected to depend on the details of the system as well as on the pulling procedure. It is important to note that, due to the simplicity of our model and the lack of experimental information on the interactions and local densities, it was not our intention to provide quantitative mapping to an exact experimental system. In fact, our simulations clearly show that when using different models that assume different protein-cortex interactions and different degrees of the membrane and linkers’ diffusivities, the value of the critical linker density can substantially change. However, our central result – the non-linear nature of the force versus linker density curve – remains unchanged.

Our simulations and experiments were carried out in a quasi-static way, which is comparable to the rate of filopodia growth [40], and allows enough time for the linkers to exclude themselves out of the tube for low and moderate linker densities. Such a pulling is however still not infinitely slow, and for high linker densities the force is not necessarily relaxed. Hence it can be expected that the critical linker density for which the force increases will be shifted to higher values for even slower pulling and longer equilibration times, both in simulations and experiments. Likewise, in fast dynamic pulling experiments the crossover might be shifted to lower linker densities, or the initial plateau of constant force might be completely absent. Interestingly, non-linear effects similar to those reported in here were observed in dynamic tether pulling experiments via hydrodynamic flows in red blood cells. The extrusion velocity of the tube was observed to depend on the extrusion force in a nonlinear fashion, which was also attributed to the dynamical feature of membrane-cytoskeleton attachments [8].

Numerous other effects can be present in live cells that contribute to the membrane-pulling force when the level of ERM proteins is changed. This can for instance include changes in cell adhesion, and changes in the cytoskeleton architecture and flow, which all might have an influence on the global tension and/or local force. However, even in the minimal system presented in our simulations it is apparent that the role of membrane-cortex attachments might be non-trivial and that their contribution cannot be necessarily considered as a “rescaled adhesion” or “effective tension”.

Finally, in simulations we observed that the linkers have a substantial effect on the stability of protrusions, preventing their diffusion and rendering them more stable against coalescence with other tubes, well in line with previous experimental observations of the increased tube coalescence upon depletion of myosin 1 proteins. We hope that our simulations and experiments will renew discussions on the mature but still unsolved question of the influence of membrane-cortex attachments on membrane tension and membrane reshaping, and stimulate the investigation of local effects and protein dynamics in the formation of membrane protrusions.

## Author contributions

A.P. performed and analysed the Monte Carlo simulations. C.V.C. developed and analysed the molecular dynamics simulations. A.Š. conceived and supervised the research. T.L., E.C. and P.B designed, performed, and analysed the membrane pulling experiments. The manuscript was written by A.P. and A.Š. All authors revised and approved the manuscript.

## Acknowledgments

We thank Ewa Paluch, Alba Diz-Muñoz, Guillaume Salbreux, Guillaume Charras, and Shiladitya Banerjee for helpful discussions. We acknowledge support from the Engineering and Physical Sciences Research Council (A.P. and A.Š.), the UCL Institute for the Physics of Living Systems (A.P., C.V.C. and A.Š.), the Royal Society (C.V.C. and A.Š.), the European Research Council (Starting grant EP/R011818/1 to A.Š.; E.C. and P.B. are partners of the advanced grant, project 339847), from Institut Curie (E.C. and P.B.) and Centre National de la Recherche Scientifique (CNRS) (E.C. and P.B.). The P.B. and E.C. groups belong to Labex CelTisPhyBio (ANR-11-LABX0038) and to Paris Sciences et Lettres (ANR-10-IDEX-0001-02). T.L. received a PhD grant from Paris Sciences et Lettres Research University and support from the Institut Curie.

## Simulation details

The membrane was modeled as a dynamically triangulated sheet with hard sphere beads situated at every vertex of a triangular lattice [41–43]. The system contains two types of beads, one type representing the membrane phospholipid patches and another corresponding to ERM (Ezrin, Radixin or Moesin) proteins. These proteins are responsible for linking the membrane phospholipids and transmembrane proteins to the underlying cytoskeletal components [16]. The dynamics of the system are governed by a Monte Carlo algorithm that involves two types of moves: vertex displacement moves and bond flip moves (Fig. S1). The vertex displacement moves mimic the lateral diffusion of lipids and proteins and allows for vertical membrane fluctuations. The bond flip moves ensure that the membrane preserves its fluidity by dynamically rearranging the connectivity. This membrane model captures correct mechanical properties of biological membranes – bending rigidity, fluidity, and the response to external forces. The model however does not describe the bilayer explicitly and the effects such as interleaflet friction are not included. The model also does not allow for topology changes such as poration or tube scission.

A tube is drawn from the membrane by applying an external point force to a predetermined lipid bead. The simulation methodology aims to mimic a slow pulling procedure such as optical tweezers or an atomic force microscopy experiment. The system’s energy is calculated using the Hamiltonian described in equation (1). This takes into account the energy required for bending a membrane patch, the energy required to create new surface area as well as the harmonic potential imposed to extrude the tube. The membrane to cortex attachment energy is controlled by a series of harmonic springs that can unbind at distances sufficiently far from the vertical equilibrium position.

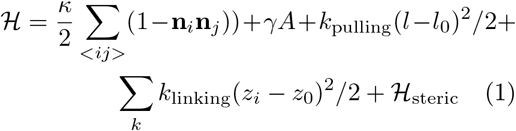

where *κ* is the membrane’s bending rigidity, i, j are pairs of triangles sharing an edge, **n**_*i*_ and **n**_*j*_ are the normal vectors of the two triangles,*γ* is the membrane surface tension and *A* is the total membrane area. *k*_pulling_ is the pulling constant that controls the tube extrusion, *l* is the vertical position of the pulled membrane bead and *l*_0_ is its target equilibrium position, which is varied between 20*σ* and 100*σ* to obtain the force-extension curves. The linkers are attached to the membrane through an harmonic potential controlled by the spring constant *k*_linking_ (Fig. 2a). The vertical position of each linker is denoted by *z*_*i*_ with *z*_0_ being its equilibrium vertical position. ℋ _steric_ is a term that accounts for the hard sphere repulsion between beads, forbidding any two beads to be closer than *σ*. The beads are not allowed to be at distances greater than 3*σ* to avoid the formation of holes and pores.

A single Monte Carlo sweep consists of *N*_vertices_ attempts at displacing a vertex and *N*_vertices_/2 bond flip attempts, where *N*_vertices_ is equal to the total number of membrane beads. In this case *N*_vertices_ = 3600 and the initial vertical side lengths *L*_*x*_ = *L*_*y*_ = 60*σ*. The acceptance probability of a Monte Carlo move is controlled by a simple Metropolis-Hastings criterion p = min[1, exp(−Δ*E/k*_*B*_*T*)], where Δ*E* is the energy change after a potential Monte Carlo move. The simulation is run in the NA_*p*_T ensemble, where A_*p*_ is the membrane’s projected area. The bending rigidity is set to *κ* = 20*k*_*B*_*T*, and the surface tension is kept constant at *γ* = 3*k*_*B*_*T/σ*^2^. The pulling constant, *k*_pulling_ = 1*k*_*B*_*T/σ*^2^, is chosen to ensure that the extrusion is slow enough such that the membrane beads can diffuse and rearrange on the timescale of the tube extrusion. The resulting force matches those in the experiments (∼ 10pN) when by taking the MC unit length to be *σ*=10 nm (which also matches the experimentally observed tube radii of tens of nm.)

The linkers are subjected to an additional harmonic potential with constant *k*_linking_ = 3*k*_*B*_*T/σ*^2^, but they are allowed to freely diffuse in the membrane’s plane. The linkers are allowed to unbind if they are displaced from their vertical equilibrium position of *z*_0_ =− 3*σ* by more than 5*σ*, but also to rebind to maintain the simulation’s detailed balance. The choice of the equilibrium position and attachment constant did not significantly affect the results. The membrane is equilibrated for 300,000 Monte Carlo steps and then the simulations are run for 2 million steps in total. During equilibration, additional volume change moves are implemented to relax the membrane surface tension. After equilibration, the spheres on the membrane’s edge are fixed in place and the box size is kept fixed during the extrusion. We checked that the system has enough excess membrane area such that the boundary conditions do not affect our results (Fig. S3).

The force profile is recorded by running a set of simulations in which a single randomly chosen central membrane bead is pulled to incrementally increasing equilibrium positions. The membrane will thus exert a restoring force on that bead. The force profile is then reconstructed from the average values of the force at every extension. This procedure allows the reconstruction of the force-extension curve obtained in a static pulling experiment. The force experienced by the protrusion’s tip decreases as the protrusion grows. The force eventually converges to a finite nonzero value as the protrusion reaches its maximum extension (Fig. S2). The measured pulling force is calculated as *F*_pulling_ = *k*_pulling_Δ*z*, where *k*_pulling_ is the pulling force constant and Δ*z* is the distance between the point being pulled and its vertical equilibrium position. As such, the force will not reach a zero value and the protrusion will grow until it reaches an extension around which it will fluctuate. We measured the force by averaging it over the course of the 1 million Monte Carlo steps, ensuring that it reached convergence. Each force-extension curve is obtained from averaging over five different random seeds, unless otherwise specified.

## References

[1] Jianwu Dai and Michael P Sheetz. Mechanical properties of neuronal growth cone membranes studied by tether formation with laser optical tweezers. Biophysical journal, 68(3):988–996, 1995.

[2] Prithu Sundd, Edgar Gutierrez, Ekaterina K Koltsova, Yoshihiro Kuwano, Satoru Fukuda, Maria K Pospieszalska, Alex Groisman, and Klaus Ley. ‘slings’ enable neutrophil rolling at high shear. Nature, 488(7411):399–403, 2012.

[3] Amin Rustom, Rainer Saffrich, Ivanka Markovic, Paul Walther, and Hans-Hermann Gerdes. Nanotubular highways for intercellular organelle transport. Science, 303(5660):1007–1010, 2004.

[4] Aurélien Roux. The physics of membrane tubes: soft templates for studying cellular membranes. Soft Matter, 9(29):6726–6736, 2013.

[5] Raktim Dasgupta and Rumiana Dimova. Inward and outward membrane tubes pulled from giant vesicles. Journal of Physics D: Applied Physics, 47(28):282001, 2014.

[6] Lin Bo and Richard E Waugh. Determination of bilayer membrane bending stiffness by tether formation from giant, thin-walled vesicles. Biophysical journal, 55(3):509–517, 1989.

[7] Jianwu Dai and Michael P Sheetz. Membrane tether formation from blebbing cells. Biophysical journal, 77(6):3363–3370, 1999.

[8] Nicolas Borghi and F Brochard-Wyart. Tether extrusion from red blood cells: integral proteins unbinding from cytoskeleton. Biophysical journal, 93(4):1369–1379, 2007.

[9] Clément Campillo, Pierre Sens, Darius Köster, Léa-Laetitia Pontani, Daniel Lévy, Patricia Bassereau, Pierre Nassoy, and Cécile Sykes. Unexpected membrane dynamics unveiled by membrane nanotube extrusion. Biophysical journal, 104(6):1248–1256, 2013.

[10] Imre Derényi, Frank Jülicher, and Jacques Prost. Formation and interaction of membrane tubes. Physical review letters, 88(23):238101, 2002.

[11] D Cuvelier, N Chiaruttini, P Bassereau, and P Nassoy. Pulling long tubes from firmly adhered vesicles. EPL (Europhysics Letters), 71(6):1015, 2005.

[12] Gerbrand Koster, Angelo Cacciuto, Imre Derényi, Daan Frenkel, and Marileen Dogterom. Force barriers for membrane tube formation. Physical review letters, 94(6):068101, 2005.

[13] Svetlana Baoukina, Siewert J Marrink, and D Peter Tieleman. Molecular structure of membrane tethers. Biophysical journal, 102(8):1866–1871, 2012.

[14] Julian Weichsel and Phillip L Geissler. The more the tubular: Dynamic bundling of actin filaments for membrane tube formation. PLoS computational biology, 12(7):e1004982, 2016.

[15] N Ramakrishnan, KK Sreeja, Arpita Roy-choudhury, David M Eckmann, Portonovo S Ayyaswamy, Tobias Baumgart, Thomas Pucadyil, Shivprasad Patil, Valerie M Weaver, and Ravi Radhakrishnan. Excess area dependent scaling behavior of nano-sized membrane tethers. Physical biology, 15(2):026002, 2018.

[16] Richard G Fehon, Andrea I McClatchey, and Anthony Bretscher. Organizing the cell cortex: the role of erm proteins. Nature reviews Molecular cell biology, 11(4):276–287, 2010.

[17] Rajalakshmi Nambiar, Russell E McConnell, and Matthew J Tyska. Myosin motor function: the ins and outs of actin-based membrane protrusions. Cellular and molecular life sciences, 67(8):1239– 1254, 2010.

[18] Drazen Raucher and Michael P Sheetz. Cell spreading and lamellipodial extension rate is regulated by membrane tension. The Journal of cell biology, 148(1):127–136, 2000.

[19] Alba Diz-Muñoz, Michael Krieg, Martin Bergert, Itziar Ibarlucea-Benitez, Daniel J Muller, Ewa Paluch, and Carl-Philipp Heisenberg. Control of directed cell migration in vivo by membrane-to-cortex attachment. PLoS biology, 8(11), 2010.

[20] Audrey Gérard, Genaro Patino-Lopez, Peter Beemiller, Rajalakshmi Nambiar, Khadija Ben-Aissa, Yin Liu, Fadi J Totah, Matthew J Tyska, Stephen Shaw, and Matthew F Krummel. Detection of rare antigen-presenting cells through t cell-intrinsic meandering motility, mediated by myo1g. Cell, 158(3):492–505, 2014.

[21] Yin Liu, Natalya V Belkina, Chung Park, Raj Nambiar, Scott M Loughhead, Genaro Patino-Lopez, Khadija Ben-Aissa, Jian-Jiang Hao, Michael J Kruhlak, Hai Qi, et al. Constitutively active ezrin increases membrane tension, slows migration, and impedes endothelial transmigration of lymphocytes in vivo in mice. Blood, The Journal of the American Society of Hematology, 119(2):445–453, 2012.

[22] Michael P Sheetz. Cell control by membrane– cytoskeleton adhesion. Nature Reviews Molecular Cell Biology, 2(5):392–396, 2001.

[23] Zheng Shi, Zachary T Graber, Tobias Baumgart, Howard A Stone, and Adam E Cohen. Cell membranes resist flow. Cell, 175(7):1769–1779, 2018.

[24] F Brochard-Wyart, Nicolas Borghi, D Cuvelier, and P Nassoy. Hydrodynamic narrowing of tubes extruded from cells. Proceedings of the National Academy of Sciences, 103(20):7660–7663, 2006.

[25] Berta Gumí-Audenis, Luca Costa, Lidia Ferrer-Tasies, Imma Ratera, Nora Ventosa, Fausto Sanz, and Marina I Giannotti. Pulling lipid tubes from supported bilayers unveils the underlying substrate contribution to the membrane mechanics. Nanoscale, 10(30):14763–14770, 2018.

[26] Sarah A Nowak and Tom Chou. Models of dynamic extraction of lipid tethers from cell membranes. Physical biology, 7(2):026002, 2010.

[27] Ayako Yamada, Alexandre Mamane, Jonathan Lee-Tin-Wah, Aurélie Di Cicco, Coline Prévost, Daniel Lévy, Jean-François Joanny, Evelyne Coudrier, and Patricia Bassereau. Catch-bond behaviour facilitates membrane tubulation by non-processive myosin 1b. Nature communications, 5(1):1–8, 2014.

[28] Ricard Alert, Jaume Casademunt, Jan Brugués, and Pierre Sens. Model for probing membrane-cortex adhesion by micropipette aspiration and fluctuation spectroscopy. Biophysical journal, 108(8):1878–1886, 2015.

[29] Drazen Raucher and Michael P Sheetz. Characteristics of a membrane reservoir buffering membrane tension. Biophysical journal, 77(4):1992– 2002, 1999.

[30] Bruno Pontes, Pascale Monzo, and Nils C Gauthier. Membrane tension: A challenging but universal physical parameter in cell biology. In Seminars in cell & developmental biology, volume 71, pages 30–41. Elsevier, 2017.

[31] Bruno T Fievet, Alexis Gautreau, Christian Roy, Laurence Del Maestro, Paul Mangeat, Daniel Louvard, and Monique Arpin. Phosphoinositide binding and phosphorylation act sequentially in the activation mechanism of ezrin. The Journal of cell biology, 164(5):653–659, 2004.

[32] Dafne Chirivino, Laurence Del Maestro, Etienne Formstecher, Philippe Hupé, Graça Raposo, Daniel Louvard, and Monique Arpin. The erm proteins interact with the hops complex to regulate the maturation of endosomes. Molecular biology of the cell, 22(3):375–385, 2011.

[33] Anagha Datar, Thomas Bornschlögl, Patricia Bassereau, Jacques Prost, and Pramod A Pullarkat. Dynamics of membrane tethers reveal novel aspects of cytoskeleton-membrane interactions in axons. Biophysical journal, 108(3):489– 497, 2015.

[34] Damien Cuvelier, Imre Derényi, Patricia Bassereau, and Pierre Nassoy. Coalescence of membrane tethers: experiments, theory, ani applications. Biophysical journal, 88(4):2714– 2726, 2005.

[35] Rajalakshmi Nambiar, Russell E McConnell, and Matthew J Tyska. Control of cell membrane tension by myosin-i. Proceedings of the National Academy of Sciences, 106(29):11972–11977, 2009.

[36] Marco Fritzsche, Richard Thorogate, and Guillaume Charras. Quantitative analysis of ezrin turnover dynamics in the actin cortex. Biophysical journal, 106(2):343–353, 2014.

[37] Alba Diz-Muñoz, Daniel A Fletcher, and Orion D Weiner. Use the force: membrane tension as an organizer of cell shape and motility. Trends in cell biology, 23(2):47–53, 2013.

[38] Martin Bergert, Sergio Lembo, Danica Milovanović, Mandy Börmel, Pierre Neveu, and Alba Diz-Muñoz. Cell surface mechanics gate stem cell differentiation. bioRxiv, page 798918, 2019.

[39] Henry De Belly, Philip H Jones, Ewa K Paluch, and Kevin J Chalut. Membrane tension mediated mechanotransduction drives fate choice in embryonic stem cells. bioRxiv, page 798959, 2019.

[40] Thomas Bornschlögl, Stéphane Romero, Christian L Vestergaard, Jean-François Joanny, Guy Tran Van Nhieu, and Patricia Bassereau. Filopodial retraction force is generated by cortical actin dynamics and controlled by reversible tethering at the tip. Proceedings of the National Academy of Sciences, 110(47):18928–18933, 2013.

[41] Yacov Kantor, Mehran Kardar, and David R Nelson. Statistical mechanics of tethered surfaces. Physical review letters, 57(7):791, 1986.

[42] J-S Ho and A Baumgärtner. Simulations of fluid self-avoiding membranes. EPL (Europhysics Letters), 12(4):295, 1990.

[43] A Baumgärtner and J-S Ho. Crumpling of fluid vesicles. Physical Review A, 41(10):5747, 1990.

