## Supplementary Material for "Influence of membrane-cortex linkers on the extrusion of membrane tubes"

This Supplementary Information provides additional details about the implementation of the membrane model and the influence of various simulation and experimental parameters on the mechanics of tube extrusion, and the experimental materials and procedures employed. Furthermore, results from a parallel molecular dynamics simulations study are presented.

### I. ADDITIONAL SIMULATION DETAILS

The membrane was modeled as a triangulated thin elastic sheet with hard sphere beads situated at every vertex. The system contains two types of beads, one type representing the membrane phospholipid patches and another corresponding to linker proteins. The dynamics of the system are evolved through a Monte Carlo scheme that involves two types of moves: vertex displacement moves and bond flip moves. The vertex displacement moves mimic the lateral diffusion of lipids and proteins, whilst also allowing for vertical membrane fluctuations. The bond flip moves ensure that the membrane preserves its fluidity by dynamically rearranging the connectivity (Fig. S1).

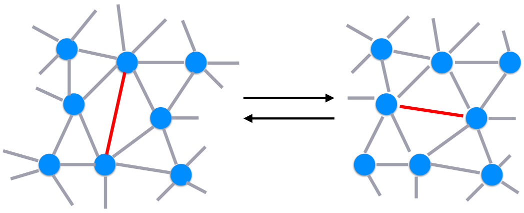

FIG. S1: **Bond flip move:** Monte Carlo moves are implemented to preserve the membrane's fluidity. The bond flips ensure that any bead will not have more than 6 neighbours at any time. These moves, together with the displacement moves, ensure that the lipids and proteins can diffuse freely in the membrane's plane.

### II. SIMULATION RESULTS

#### A. Tube pulling force relaxation profile

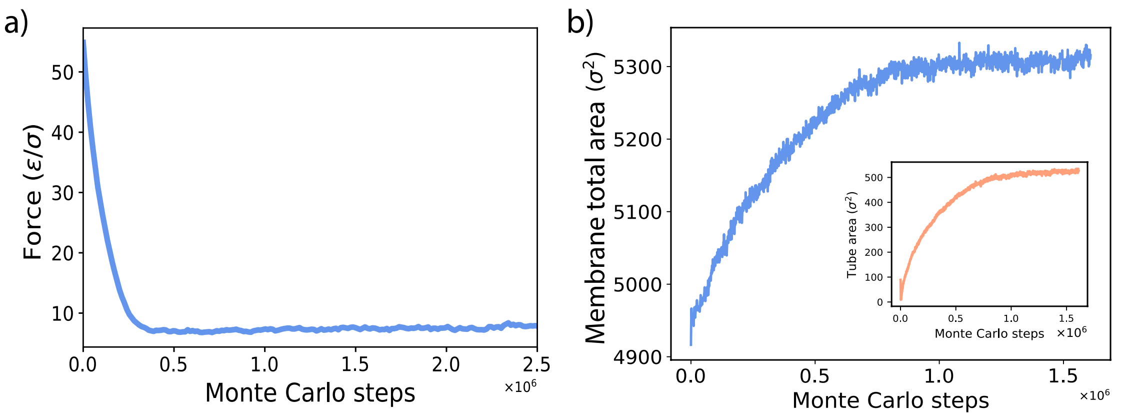


FIG. S2: **Force profile of a typical extrusion experiment.** a) The force acting on the protrusion's tip decays as the simulation proceeds, eventually converging at a non-zero value. b) The membrane area increases until it also reaches a steady state value (inset shows the tube area which follows a similar behaviour).

The point force acting on a protrusion's tip decreases until it reaches convergence at a non-zero value (Fig. S2a). The total membrane area and the tube area show a gradual increase until they stabilise at a steady state value (Fig. S2b). The force cited in the main text is calculated from an average over one million Monte Carlo steps between 1 million and 2 million Monte Carlo steps. The force-extension curves were averaged over 5 different simulation seeds each, unless otherwise mentioned.

#### B. Influence of membrane size on the measured force

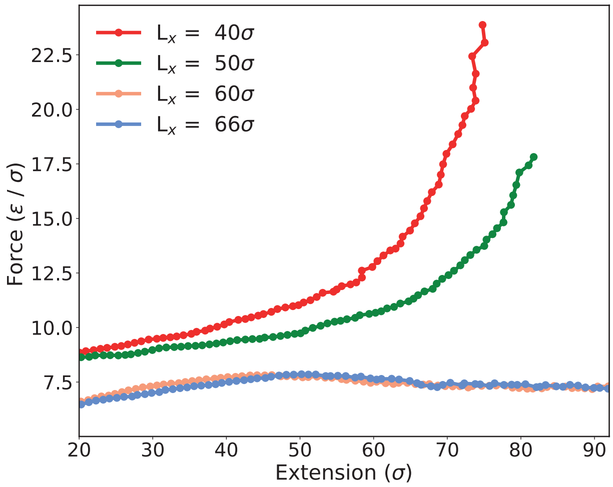


FIG. S3: **Influence of the fixed boundaries on the measured force.** The measured force increases significantly if the box size is too small, but it does not vary for large membrane sizes. The simulations presented in the main text used $L_x$=60$\sigma$.

In order to check if the finite boundaries influence the measured force, we ran simulations in which the membrane size was varied between 40$\sigma$ and 66$\sigma$. For small side lengths, we observe a clear increase in the recorded force, as the material available for tube growth is depleted leading to an increase in the tension as illustrated in Fig.S3. However, for large membrane side lengths such as the one used in simulations, this effect is not observed, indicating that there is enough excess membrane area available for tube growth. As such, the measured force is not influenced by the fixed boundaries for the system sizes presented here.

#### C. Influence of the linker proteins attachment energy on tube shape and composition

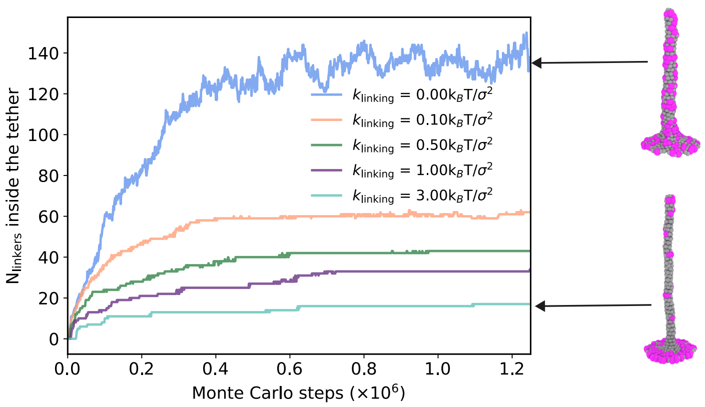


FIG. S4: **Number of linkers inside the tube for different attachment energies**. The flow of linkers inside the protein is heavily influenced by the protein binding. At higher attachment energies, the linkers will avoid the tube and partition preferentially in the membrane.

We next examined how the protein binding strength influences the tube shape and protein partitioning between the membrane and the tube. The linker attachment constant, $k_{linking}$, was varied between 0 $k_BT/\sigma^2$ and 3 $k_BT/\sigma^2$, whilst keeping the number density of proteins constant at 20%. At high linking attachment energies, the proteins partitioned preferentially into the membrane and avoided the tube entirely. This led to the formation of very thin elongated tubes with few proteins trapped inside (Fig. S4). At lower attachment energies, more linkers partitioned inside the tube, as the detachment cost is easier to overcome. The attachment energy highly influences the tube radius. The tube diameter decreases inversely proportional to the attachment energy (Fig. S5).

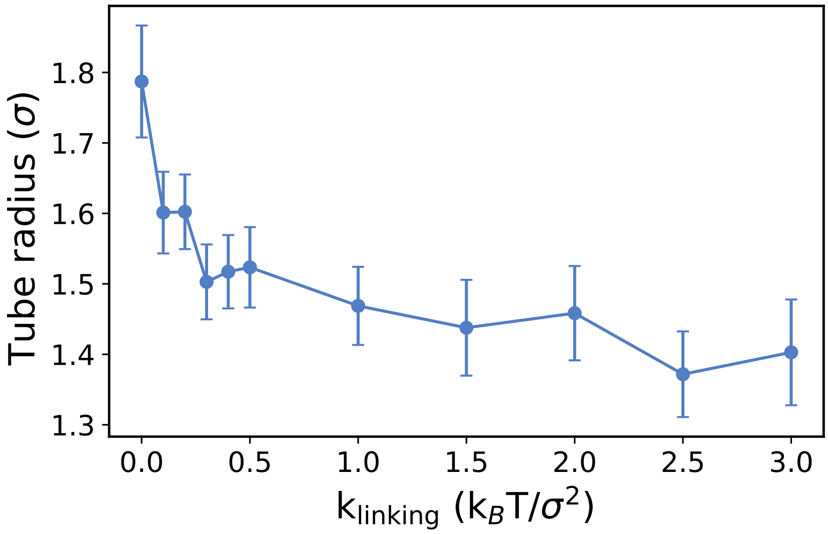


FIG. S5: **Influence of the attachment energy on tube radius**. The extruded tubes are thinner at higher attachment energies, as the proteins are pinned stronger to the membrane and hinder the free flow of lipid material inside the tube.

#### D. Influence of attachment energy on the pulling force

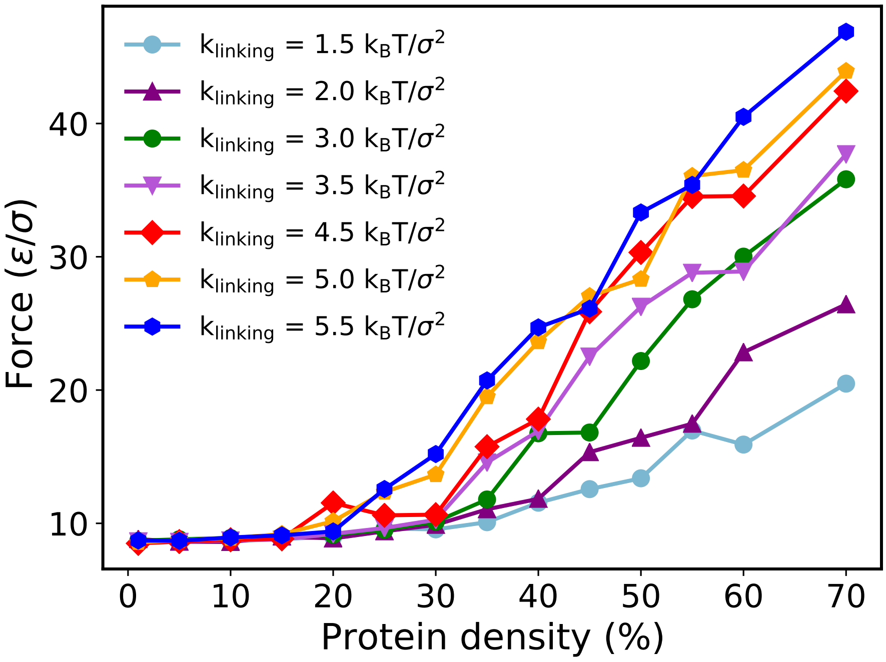


FIG. S6: **Influence of attachment energy on the shape of the force vs protein density curve.** The forces were measured for a target extension of $60\sigma$. The results are averaged over five simulation seeds.

The linkers' attachment energy also influences the values of the recorded pulling force. We studied how the critical density of linkers at which the extrusion force increases dramatically (as illustrated in Fig. 2c of the main text) is affected by the binding strength. We ran simulations in which varied the linkers' attachment constant, $k_{linking}$ between 1.5 $k_BT/\sigma^2$ and 5.5 $k_BT/\sigma^2$ for a target extension of $60\sigma$. As illustrated in S6, we observe that cross-over point from a flat to a rapid increase in the measured force can be shifted to lower protein densities by increasing the linkers' binding strength.

#### E. Distribution of linkers around the tube at different protein expression levels

The protein expression levels greatly influence the distribution profile of the linkers with respect to the tube. This, in turn, has an impact on the force measured in experiments. At low densities, the proteins reorganise around the tube and prefer segregating into protein rich areas (Fig. 3 of the main text and Fig. S7). In this regime, the protein levels do not significantly affect the force recorded in experiments (Fig. S2b). At higher densities, the proteins cannot avoid entering the tube as can be seen in Fig. S7. The linkers will be distributed uniformly across the entire membrane patch, with the tube's composition being identical to that of the membrane. The tube extrusion is thus energetically costly as it requires the detachment of the linkers from their rest positions. This leads to a dramatic increase in the measured force if the linkers are overexpressed.

#### F. Force-extension curve in the overexpression regime

The protein expression levels regulate the shape of the force-extension curve. At low protein densities, the shape of the curve is not significantly influenced by the protein density, showing a consistent characteristic force barrier followed by a flat plateau over a large extension (Fig. 2b of the main text). If the protein is overexpressed, the measured force increases dramatically as the proteins will be forced to detach from the membrane and they will be included in the tube (Fig. S8). The measured force will

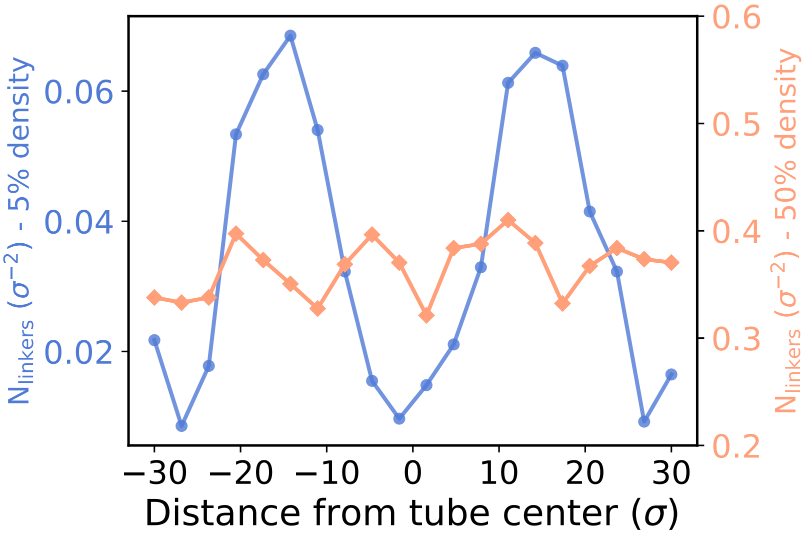


FIG. S7: **Variation of the number of linkers per unit area with the distance from the tube**. The number of linkers per unit of membrane area at 5% and 50% protein density.

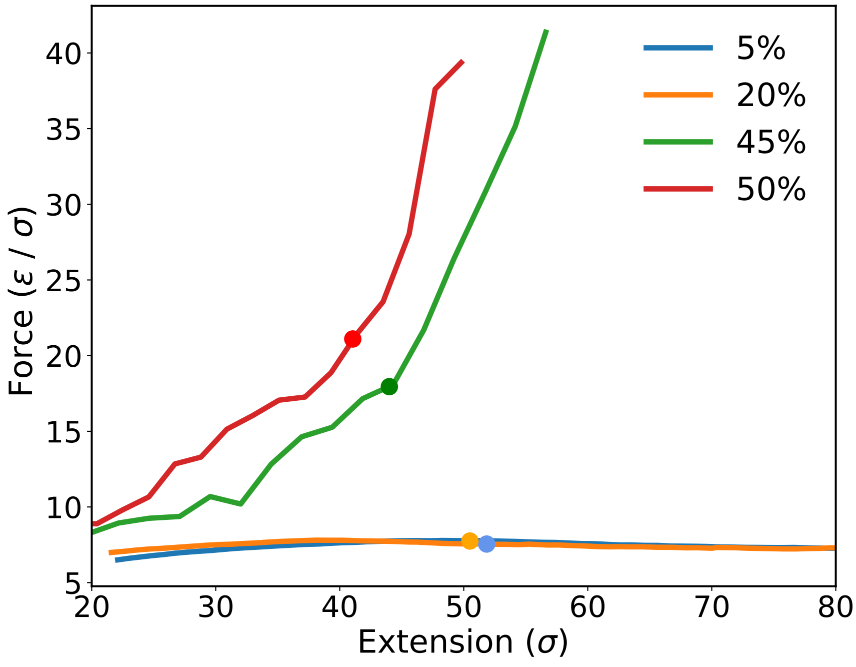


FIG. S8: **Force-extension curves at low and high linker densities**. At low densities, the force does not vary significantly over a wide range of tether extensions. The force increases substantially if the protein is overexpressed due to the high energetic cost associated with detaching the linkers from their rest positions. The markers correspond to the force and extension values recorded for a target extension of $60\sigma$, as quoted in Fig. 2c of the main text.

thus be significantly greater in this linker expression regime (as can also be observed from Fig. 2c of the main text).

#### G. Comparison with non-diffusive linkers

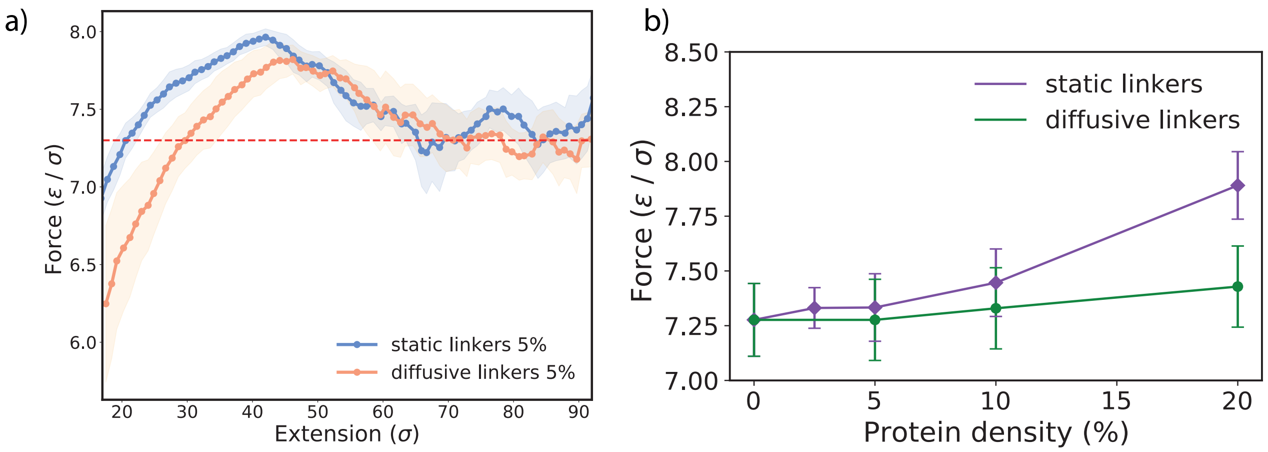


FIG. S9: **Force measurements for static linkers**. a) Comparison between the force-extension curves of static and diffusive linkers at 5% protein density. b) Force vs. protein density at a target extension of $80\sigma$ for both the static and diffusive linkers.

In order to check how the diffusivity of the linkers influences the results presented in the main text, we ran simulations in which the membrane linkers are not allowed to diffuse laterally for more than $0.3\sigma$ from their post-equilibration position during the tube extrusion. The linkers are still allowed to detach vertically and to rebind as previously described. At small linker densities, the static linkers display a slightly higher barrier to extrude a tube, but once the tube has undergone the shape transition, the plateau force required to pull the tube is identical to one for diffusive linkers as illustrated in Fig. S9. This is because the membrane flows around the linkers and enters the tube, while the linkers still avoid entering the tube. At higher expression levels, in the case of the static linkers, the plateau force increases as the membrane redistribution around them is not possible any more and the linkers need to detach for the tube to grow. This result is qualitatively similar to the one from the dynamic linkers. The only difference in the force-extension between the two is that the cross-over between the two regimes occurs at lower linker densities in the case of static linkers, as shown in Figure S9b. In addition, in the case of static linkers, the linker clustering is not possible and hence not observed.

#### H. Molecular dynamics simulations

Throughout this work we have investigated the role of membrane-cortex linkers on the extrusion of membrane tubes using Monte Carlo simulations of planar lipid sheets. To further support our observations we complement these results with molecular dynamics simulations of large spherical lipid vesicles subject to an extruding force in the presence of linker proteins. In this section we will discuss the details and implementation of such computer simulations and present some of the most relevant results, discussing the implications for the membrane tether extrusion process.

The system under study consists of an initially relaxed sphere of $N_{\text{particles}} = 48002$ membrane and protein particles giving a radius of $R \sim 56\ \sigma$. The vesicle contains two types of beads, one representing the membrane lipids (coloured in blue in Fig. S8) and another corresponding to ERM (Ezrin, Radixin or Moesin) proteins (coloured in green in Fig. S8). Both are modelled using the single particle thick model developed by Yuan et al. [1] which reproduces biologically relevant mechanical properties of membranes. The specific parameters of the model are chosen to encode for a membrane of vanishing spontaneous curvature and bending rigidity of 15 $k_BT$, $k_B$ being the Boltzmann constant and $T$ being temperature. Using the original paper's notation: $\epsilon = 4.34\ k_BT$, $\zeta = 4$, $\mu = 3$, $r_{\min} = 1.12\ \sigma$, $r_c = 2.6\ \sigma$

and $\theta_0 = 0$. The choice of the pulling force again matches those measured in experiments ( 10pN) when taking $\sigma = 10nm$. It is important to note that, unlike the triangulated membrane model, this membrane model allows for membrane poration and breakage. Hence for substantially higher pulling forces the would break.

To account for the membrane-cortex cross-linking of the ERM beads, we introduce an additional ghost particle (coloured in yellow in Fig. S8, not subject to any dynamics) at the sphere's centre, tethering the linker beads to an equilibrium distance $r_{\rm link} = 54\ \sigma$ via a harmonic potential of constant $k_{\rm link} = 1\ k_BT\ /\sigma^2$. It should be noted that this potential only imposes a preferred radial distance to the vesicle's centre, but it allows the protein beads to freely diffuse on the membrane surface. Finally, we model the tube extrusion process via an external bead (coloured in cyan in Fig. S8) of size $r = 3\ \sigma$ subject to a harmonic potential of constant $k_{\rm pull}$ pulling it to an equilibrium target extension $Z$ above its initial position. The particle is initially positioned above the vesicle and is bound to it via a Lennard-Jones potential of strength $\epsilon = 6\ k_BT$, with an equilibrium distance $r_{min} = 2\ \sigma$ and cutoff at $r_c = 5\ \sigma$. The system's conditions are therefore fully defined by only two free parameters: the ERM protein density, which we vary between 0% and 10%, and the target extension $Z$ (the pulling potential's contant $k_{\rm pull}$ is adjusted for an initial pulling force $F_0 = k_{\rm pull}\ Z = 30\ k_BT\ \sigma^{-1}$) which we vary between $Z = 10\ \sigma$ and $Z = 400\ \sigma$. We run simulations using the open source molecular dynamics package LAMMPS [2] with a Langevin thermostat (at $T = 1\ \epsilon/k_B$ and damping coefficient of 1) within the NVE ensemble for $200,000$ time-steps, each of size 0.01 simulation time units ($\sqrt{m\ \sigma^2\ \epsilon^{-1}}$), after and initial 2000 relaxation steps (without pulling) to allow for equilibration. All results presented in this section are the result of three independent runs for each simulation setup.

The first notable result we recover via the molecular dynamics simulations, despite the significant geometry differences with respect to the Monte Carlo simulations discussed in the main text, is the redistribution of ERM proteins as a result of the tube extrusion. As displayed in Fig. S10, protein density fluctuates around its average value far from the tube but rapidly increases as we approach the tube's base until reaching a peak below the extruded tube (note how the maximum is consistently situated around $z = 52.5\ \sigma$ for all three curves in Fig. S10) before rapidly dropping to zero at the base and inside the tube. Indeed, consistent with our previous results (see Fig. 3 in the main text), we observe again that linkers are excluded from the tube and accumulate around it immediately following extrusion.

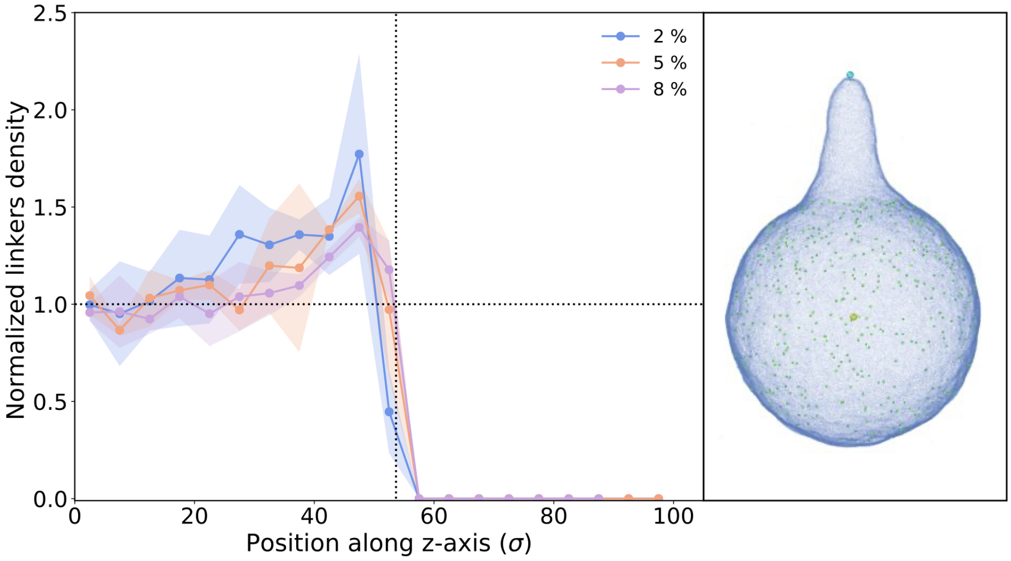


FIG. S10: **Linkers are excluded from the extruded tube, resulting in an accumulation in the periphery of the tube's base.** The local density of proteins along the z-axis (only positive direction) normalized by the average density of the particular setup (2, 5 and 8 % respectively, see legend). Each point represents the protein density on a membrane slice of width $w = 5\sigma$ centered at a certain height $z$ above the midpoint of the vesicle, centred over three independent runs. Shaded regions correspond to the standard deviation of measurements. The vertical dotted line indicates the tube base position. The snapshot to the right serves as an example to illustrate this accumulation phenomenon for the 2 % case. All three different setups correspond to a target extension of $Z = 100\ \sigma$.

Furthermore, another feature of the tube extrusion process we reproduce via molecular dynamics simulations is the dependency of the tube radius on the tube's extension and the protein density (see Fig. 2d in the main text). As shown in Fig. S11, the tube radius exhibits a similar decaying profile with tube extension for different protein densities, manifesting the transition from a cone-like structure to a thin elongated tube, as displayed in the two insets. Furthermore, we also observe how, for a given tube extension, the linker density has a modulating effect on the tube's radius which narrows as the density increases (displayed in the three snapshots to the right of Fig. S11). As discussed in the main text, this effect is a direct result of how an increase in ERM linkers limits the amount of freely available membrane to be extruded (see Fig. 2d and corresponding discussion in the main text).

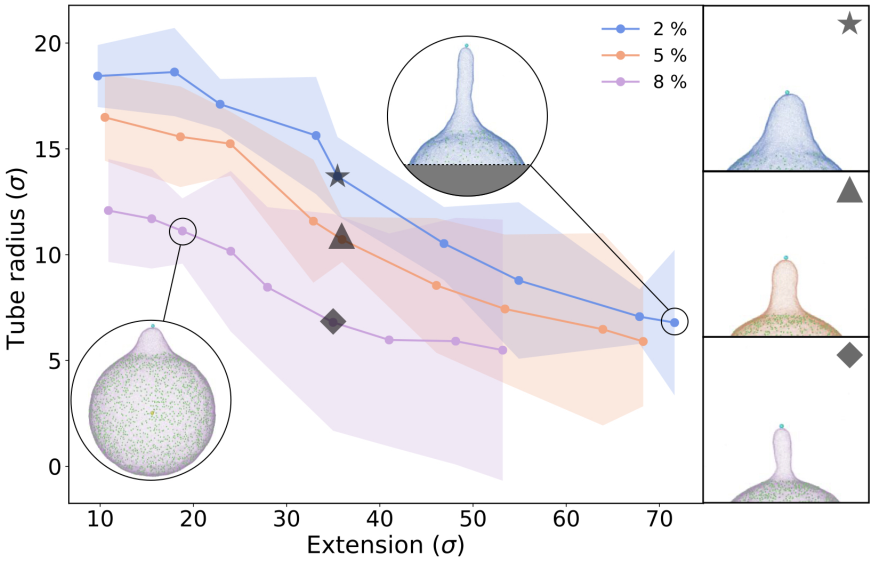


FIG. S11: **Radius of extruded tubes decreases with tube height and protein density.** Radius of the extruded tube measured at different equilibrium extensions for three ERM protein density (colour-coded, see legend). Each point is the result of averaging over three independent runs. Shaded regions correspond to the standard deviation of measurements. Insets and snapshots to the right exemplify the equilibrium situation for different setups (see symbols). Lipid beads are coloured according to protein density following the legend.

Finally, we also reproduce the strong increase in equilibrium pulling force for high protein densities (see Fig. 2c and corresponding discussion in the main text). As presented in Fig. S12 , we observe that in the lower range of explored linker densities the equilibrium force (computed as $F = k_{\text{pull}}\,\Delta z$, see main text) for a given target extension remains fairly low and constant and does not display any noticeable changes as the density increases into intermediate ranges. However, as we reach the higher values of ERM protein densities (i.e. 9% and 10%) we observe a sharp increase in the equilibrium pulling force (close to or even above double the low density values).

In summary, by implementing an alternative computational approach to investigating the role of membrane-cortex linkers on the extrusion of membrane tubes, we are able to provide evidence of the universality of some of the main features discussed in the main text of this work. Indeed, in this section we have shown that key elements of this process such as protein redistribution, tube geometry modulation by protein density and tube elongation or even the substantial increase in the required force for higher protein concentrations are all present independently of the membrane geometry and the exact simulation model.

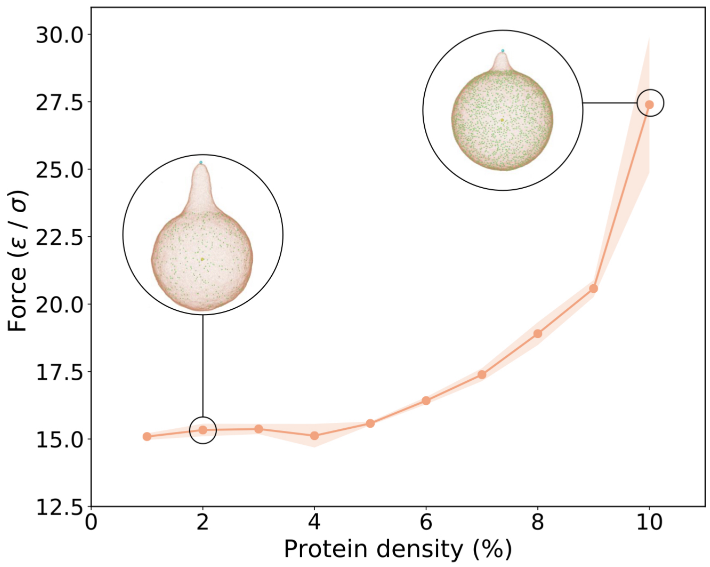


FIG. S12: **Force on the pulling bead increases substantially for high protein densities.** Equilibrium pulling force, averaged over three independent runs for each setup, measured at different ERM protein densities for target extension $Z = 100\sigma$. The shaded region corresponds to the standard deviation of measurements. The insets exemplify the equilibrium configuration of the system under different conditions.

### III. EXPERIMENTAL RESULTS

#### A. Tether force convergence

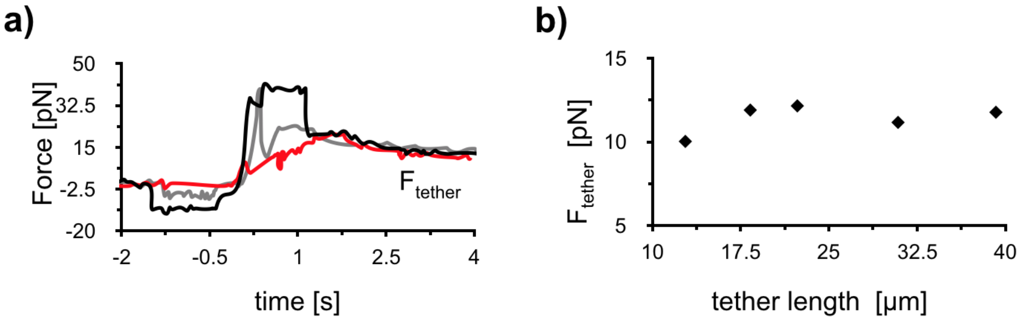


FIG. S13: **Tether forces rapidly converge to $\mathbf{F}_{\text{tether}}$ and are independent of the tether length.** a) Full force profile corresponding to three membrane tethers pulled from different cells during one experiment. While the initial force profile is very different for each of the cells since its depends on the contact area between the bead and the cell membrane, individual tether holding forces converge to similar values within seconds. b) Tether holding force of an individual membrane tether as a function of tether length.

### B. Tube pulling force for different cell types

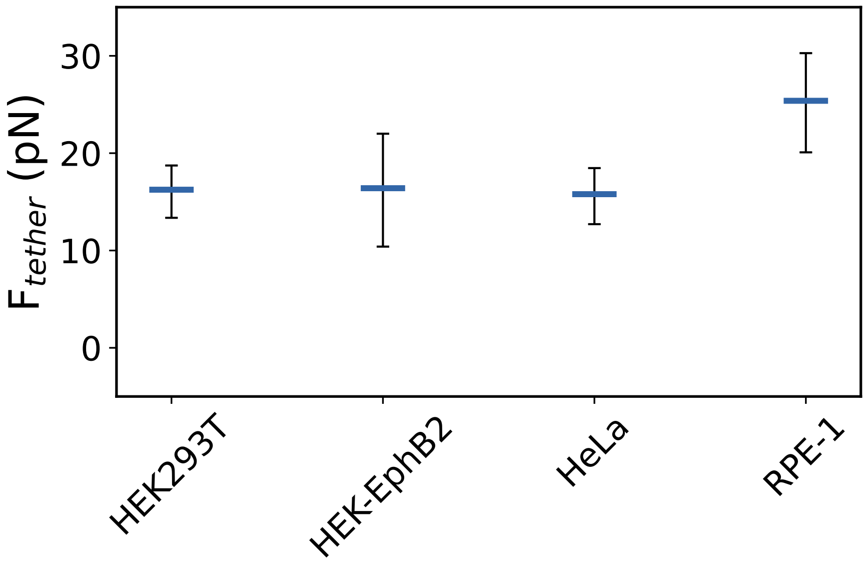


FIG. S14: **Static tether forces depend on cell type.** Tether forces in HEK293T (n=5), HEK-EphB2 (n=16), HeLa (n=5) and RPE-1 (n=5) cells.

### C. Expression levels of proteins analyzed by western blot and immunofluorescence

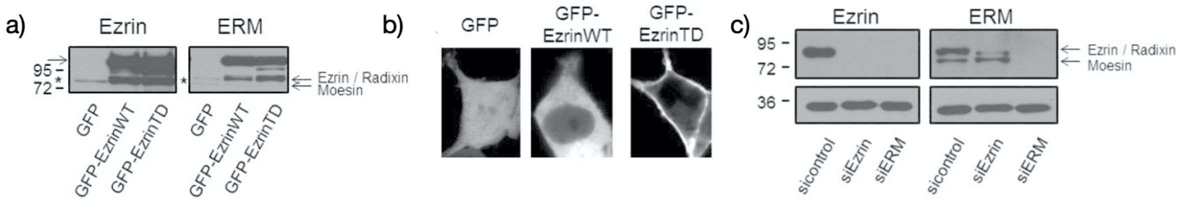


FIG. S15: **Expression levels of proteins analyzed by western blot and immunofluorescence.** (a) Western Blots showing expression levels of proteins after expressing GFP labeled constructs in HEK293T cells. Expression level of GFP-Ezrin is superior to the endogenous ezrin (*) and ERM. (b) Representative equatorial plane fluorescence images of HEK293T cells expressing fluorescent recombinant proteins. Note that we selected cells with similar expression levels of fluorescent proteins. GFP-EzrinWT is detected in the cytoplasm and at the plasma membrane and GFP-EzrinTD is predominantly at the plasma membrane. (c) Western blots showing expression levels of proteins after depletion using siRNA, detected with antibodies against Ezrin and ERM, respectively. All siRNA treatments reduce the protein levels below detection levels. GAPDH is the loading control. Note that Ezrin and Radixin are not well separated and correspond to one band (upper band) when detected with anti-ERM antibodies. Thus this band seems slightly shifted and only partially reduced in cells treated by siEzrin. The 3 ERM are fully depleted with siERM.

#### D. Fitting of relaxation curves

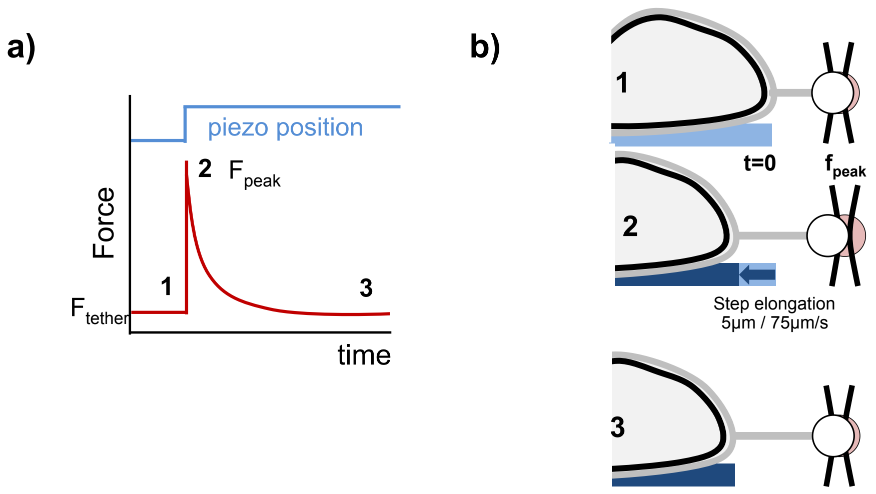


FIG. S16: **Step elongation of existing membrane tethers to assess tether relaxation.** a) Three force regimes can be observed during step elongation of an existing membrane tether. b) A relaxed membrane tether with holding force $F_{tether}$ (1) is elongated by imposing a fast step displacement of a piezo-controlled stage leading to a peak in force (2). Over time the force relaxes back to the initial value of the tether force ($F_{tether}$).

A 5 $\mu$m extension at 75 $\mu$m/s was applied on a preformed tube held at a static tether force $F_{tether}$ (Fig. S13). During elongation we observed a rise up to a peak force $F_{peak}$ when elongation is stopped, defined as $t = 0$. We measured the force relaxation over time $F(t)$ at constant tether length in the presence or in the absence of ERM (Fig. 5b of the main text). Early work on tether pulling from cells generally considered a single exponential relaxation for the force [3–5]. Recent work both on vesicles with a reconstituted actin cortex and cellular membranes vesicles revealed that the force relaxation cannot satisfyingly be described with a single exponential, but rather contains both a fast exponential decay (on the order of seconds) independent of actin and a linear slow decay that depend on actin and appears diffusive [6]. We confirmed that a single exponential decay does not account properly for our data. In contrast, our data is correctly described by a simple model of a composition gradient across the neck: the tension gradient, reflected by the tether force, is dissipated with a characteristic relaxation time that depends on membrane-component diffusion and thus possibly on the density of actin-PM linkers that form 'obstacles', thus reducing their diffusion. We have fitted the force relaxation data (Fig. 5b of the main text) with:

$$F(t) = \sqrt{F_{\text{peak}}^2 - (F_{\text{peak}}^2 - F_0^2)g(t/\tau)} \tag{S1}$$

where:

$$g(t/\tau) = 1 - e^{-(t/\tau)}(\text{erf}(c\sqrt{t/\tau})) \tag{S2}$$

with the relaxation time $\tau$ as the only free parameter. $\tau$ is expected to depend on the diffusion of the minority component in the cell membrane, which in turn is expected to be influenced by the density of linkers at the interface between plasma membrane. Our data set fits well with equation S1 (Fig. 5b in the main text). As expected, the relaxation time is reduced when ERM are depleted (0.7 s in the control and 0.3 s after ERM depletion, Fig. 5b).

### IV. MATERIALS & METHODS

**Cell culture.** HEK293T, HEK-EphB2 [7], HeLa, RPE-1 and NIH3T3 cells were cultured at 37°C and 5% CO2 in DMEM GlutaMAX medium (gibco 61965-026), supplemented with 10% fetal bovine serum (gibco 10270106), and 1% Penicillin Streptomycin (gibco 15070063).

**Plasmids**
Carboxy-terminally-tagged GFP-EzrinWT and and GFP-EzrinTD [8, 9] fusion protein were previously described.

**siRNA.** siEzrin (L-017370-00), ERM triple depletion as described in [10] with siEzrin (EZR J-017370-08), siRadixin (RDX J-011762-07), siMoesin (MSN J-011732-08), and non-targeting control siRNA (D-001810-10), all from Dharmacon.

**Transfection.** Transfection of cells with protein-encoding plasmids was performed using Lipofectamine$^{TM}$ 2000 (Invitrogen 11668-019). 250,000 cells per well were seeded in a 6 well plate (TPP 92006) 24 hours prior transfection with 1 $\mu$g of the other plasmids, per well. The transfection mix was prepared as follows: 5 $\mu$L of Lipofectamine$^{TM}$ 2000 and the respective amount of DNA were added to 250 $\mu$L of OptiMEM (gibco 31985-062) each in a separate 2 mL microtube (Sarstedt 72.695.500). After 5 minutes of incubation at room temperature, both solutions were united and mixed by pipetting up and down several times and left to incubate at room temperature for 15 minutes. The resulting 500 $\mu$L of transfection mix was added to the corresponding well, containing 1.5 mL freshly added, pre-warmed full medium. The medium was exchanged for full medium 6-8 hours after transfection and cells were assessed after ∼24h of transfection.

Cells were transfected with siRNA using Lipofectamine$^{TM}$ RNAiMAX (Invitrogen 13778-030). The transfection mix was prepared as follows: 5 $\mu$L of Lipofectamine$^{TM}$ RNAiMAX and the respective amount of siRNA were added to 250 $\mu$L of OptiMEM (gibco 31985-062) each in a separate 2 mL microtube (Sarstedt 72.695.500). After 5 minutes of incubation at room temperature, both solutions were united and mixed by pipetting up and down several times and left to incubate at room temperature for 30 minutes. The resulting 500 $\mu$L of transfection mix was added to the corresponding well, containing 1.5 mL freshly added, pre-warmed full medium.

For depletion of Ezrin, ERM, and respective controls, 80,000 cells were seeded out 24h prior treatment and transfected twice, the secondary transfection following 48 hours after the first. Cells were assessed after a total of ∼96 hours. The respective concentrations of siRNA were 20 nM for Ezrin, and 10 nM for each siRNA of Ezrin, Radixin, and Moesin, thus yielding a total of 30 nM for ERM depletion.

**Western blots.** After washing with PBS cells were lysed in 1.5x SDS-loading buffer supplemented with $\beta$-mercapto-ethanol pre-heated to 95°C. After treatment with (Sigma),benzonase cell lysates were analyzed by SDS-PAGE and transferred to nitrocellulose membranes. Ezrin and ERM were detected with antibodies indicated on the figures.

**Experimental chamber.** Cells were grown on 25 mm circular glass cover slips (VWR ECN 631-1584) prepared in the following way: 1) Cleaning with Isopropanol, Ethanol, water, Ethanol and then dried under a cell culture hood. 2) Coating with a 20 $\mu$g/mL solution of Laminin (Sigma L2020) for ∼2 hours at 37°C. Cells were analyzed 16-20 hours after seeding out 250000 cells per glass coverslip. The medium was exchanged for CO2-independent DMEM medium (gibco 21063-029), supplemented with 1.5 mg/mL $\beta$-Casein (Sigma C6905) to passivate the surface (experiment medium), 1 hour before a measurement. The glass cover slips were fixed on a custom-made experimental chamber using vacuum grease (Sigma Z273554). The cells were maintained in experimental medium throughout the experiment supplemented with carboxylated polystyrene beads (Spherotech CP-30-10, 3.07 $\mu$m nominal diameter, 0.001 % suspension) used for tether pulling. The temperature was maintained at ∼37°C throughout the experiment using a custom-built objective-heater based on a resistive collar.

**Fluorescence images.** Transfected cells were imaged using a conventional confocal microscope at the equatorial plane to assess expression and localization of fluorescently tagged proteins prior to tube pulling. In our measurements, besides excluding GFP-negative cells and those showing clear morphological aberrations after over-expression, we kept the same imaging conditions for our experiments. The same laser power and PMT settings were used across different overexpression constructs and only cells within the dynamic range were selected to perform experiments.

**Optical Tweezer.** The custom-built optical tweezer set-up consisted of a 1064 nm Ytterbium fiber laser (IPG Photonics) and a Nikon C1 Plus confocal microscope. We employed a Nikon CFI

Plan Fluor 100x, NA 1.3 oil immersion objective which was surrounded by a resistive collar to allow heating of the objective to maintain a stable temperature of 37°C within the experimental chamber. The sample chamber was fixed on a voltage-controlled piezo-drive (MadCityLabs Nano-LP100). Bright field images of the bead were acquired using an EM-CCD camera at a frame rate of ∼180 frames per second (iXon DU-897, Andor), and analyzed by a custom MATLAB script. Forces were calculated from bead positions following the equation: $F = k(x - x_0)$, where $k$ is the trap stiffness, $x$ the position of the bead, and $x_0$ the reference position at equilibrium. Trap stiffness calibration was obtained using the viscous drag method, including Faxen's correction for calibration close to surfaces [11], with a trap stiffness adjusted to 44 pN/$\mu$m at 3W input power (400mW at the back focal plane of the objective).

**Tube pulling.** Using a custom LabVIEW program to control the piezo stage, tubes were pulled, by trapping an isolated floating bead, bringing it into contact with the cell for a short period (∼3 seconds), and then moving the cell away from the trap center in x direction using the piezo. For measuring the static tether force, the tube was about 10$\mu$m long and held steady for at least 10 seconds. For the dynamical force measurements, the initial tether had a length of about 10$\mu$m. While the initial force overshoot for tether formation varied widely, a relaxation of the force to a stable plateau could generally be observed within the first 10 seconds after tube formation.

**Step elongation.** Once the tether holding force was stabilized, step elongation of the tube was performed by imposing a step-displacement of the piezo stage away from the trap center of 5 $\mu$m at a speed of 75 $\mu$m/s. The force relaxation was then recorded.
The change in force is monitored by imaging the bead holding the tube at high temporal resolution (175 fps). This allows following the peak in force upon step elongation and the subsequent relaxation.

**Statistics.** Data sets for effective membrane tension and relaxation timescale were analyzed by one-way ANOVA. If the null hypothesis was successfully rejected, F-tests were conducted on a paired basis to compare variances and, depending on the outcome, two-tailed t-tests assuming equal or unequal variances were employed. Only p-values smaller than 0.05 are displayed in the figures.

---


[1] Hongyan Yuan, Changjin Huang, Ju Li, George Lykotrafitis, and Sulin Zhang. One-particle-thick, solvent-free, coarse-grained model for biological and biomimetic fluid membranes. *Phys. Rev. E*, 82:011905, Jul 2010.

[2] Steve Plimpton. Fast parallel algorithms for short-range molecular dynamics. *Journal of computational physics*, 117(1):1–19, 1995.

[3] William C Hwang and Richard E Waugh. Energy of dissociation of lipid bilayer from the membrane skeleton of red blood cells. *Biophysical Journal*, 72(6):2669, 1997.

[4] Liselotte Jauffred, Thomas Hønger Callisen, and Lene Broeng Oddershede. Visco-elastic membrane tethers extracted from escherichia coli by optical tweezers. *Biophysical journal*, 93(11):4068–4075, 2007.

[5] Zhiwei Li, Bahman Anvari, Masayoshi Takashima, Peter Brecht, Jorge H Torres, and William E Brownell. Membrane tether formation from outer hair cells with optical tweezers. *Biophysical journal*, 82(3):1386–1395, 2002.

[6] Clément Campillo, Pierre Sens, Darius Köster, Léa-Laetitia Pontani, Daniel Lévy, Patricia Bassereau, Pierre Nassoy, and Cécile Sykes. Unexpected membrane dynamics unveiled by membrane nanotube extrusion. *Biophysical journal*, 104(6):1248–1256, 2013.

[7] Marie-Thérèse Prospéri, Priscilla Lépine, Florent Dingli, Perrine Paul-Gilloteaux, René Martin, Damarys Loew, Hans-Joachim Knölker, and Evelyne Coudrier. Myosin 1b functions as an effector of ephb signaling to control cell repulsion. *Journal of Cell Biology*, 210(2):347–361, 2015.

[8] Richard F Lamb, Bradford W Ozanne, Christian Roy, Lynn McGarry, Christopher Stipp, Paul Mangeat, and Daniel G Jay. Essential functions of ezrin in maintenance of cell shape and lamellipodial extension in normal and transformed fibroblasts. *Current Biology*, 7(9):682–688, 1997.

[9] Sylvie Coscoy, François Waharte, Alexis Gautreau, Marianne Martin, Daniel Louvard, Paul Mangeat, Monique Arpin, and François Amblard. Molecular analysis of microscopic ezrin dynamics by two-photon frap. *Proceedings of the National Academy of Sciences*, 99(20):12813–12818, 2002.

[10] Dafne Chirivino, Laurence Del Maestro, Etienne Formstecher, Philippe Hupé, Graça Raposo, Daniel Louvard, and Monique Arpin. The erm proteins interact with the hops complex to regulate the maturation of endosomes. *Molecular biology of the cell*, 22(3):375–385, 2011.

[11] Keir C Neuman and Steven M Block. Optical trapping. *Review of scientific instruments*, 75(9):2787–2809, 2004.
